# Probing the molecular mechanism of the action of *Deinococcus radiodurans* RecD2 at single-molecule resolution

**DOI:** 10.1101/2021.12.21.473662

**Authors:** Debayan Purkait, Farhana Islam, Padmaja P. Mishra

## Abstract

Helicases are ATP-driven molecular machines that directionally remodel nucleic acid polymers in all three domains of life. Helicases are responsible for resolving double-stranded DNA (dsDNA) into separate single-strands and this activity is essential for DNA replication, nucleotide excision repair, and homologous recombination. RecD2 from *Deinococcus radiodurans* (DrRecD2) has important contributions towards the organism’s unusually high tolerance to gamma radiation and hydrogen peroxide. Although previous X-ray Crystallography studies have revealed the structural characteristics of the protein, the direct experimental evidence regarding the dynamics of the DNA unwinding process by DrRecD2 in the context of other accessory proteins is yet to be found. In this study, we have probed the exact binding event and processivity of DrRecD2 at single-molecule resolution using Protein-induced fluorescence enhancement (smPIFE) and Forster resonance energy transfer (smFRET). We have found that the protein prefers to bind at the 5 ‘ terminal end of the single-stranded DNA (ssDNA) by Drift and has helicase activity even in absence of ATP. However, a faster and iterative mode of DNA unwinding was evident in presence of ATP. The rate of translocation of the protein was found to be slower on dsDNA compared to ssDNA. We also showed that DrRecD2 is recruited at the binding site by the single-strand binding protein (SSB) and during the unwinding, it can displace RecA from ssDNA.

## Introduction

Helicases are ATP-driven molecular machines that directionally remodel nucleic acid polymers in all three domains of life. Helicases are responsible for resolving double-stranded DNA into separate single-strands and this activity is essential for DNA replication, nucleotide excision repair, and homologous recombination[1–3]. Following their various functions in different cellular pathways, these proteins show tremendous diversity in the context of their structures and mechanisms of action and thus have been classified into six superfamilies (SFs)[4]. Superfamily 1 (SF1) is composed of helicases that primarily function in DNA repair pathways, members of this family include *Escherichia coli* UvrD, Rep helicases, and RecD. SF1 is further subdivided into SF1A and SF1B based on the directionality of translocation along with the nucleic acid polymer. SF1A helicases translocate from the 3’ to 5’ end of the double-stranded DNA and SF1B helicases, like RecD and RecD2, perform the task in reverse direction[5].

The bacterial RecD (Recombinase D) helicase family is represented by RecD (or RecD1) and RecD2[6]. RecD2 helicase is typically found in bacterial systems that lack RecBCD complex, which has been well studied for decades and is now a standard molecular model system for many textbooks of genetics. In *E. coli*, the double-stranded DNA breaks (DSBs), the most lethal outcome of ionising radiations and other DNA-damaging agents, are repaired by the RecA-dependent RecBCD homologous recombination pathway. The RecBCD heterotrimeric complex consists of three individual units of RecB (SF1A helicase), RecC (nuclease), and RecD, binds to the dsDNA as a whole and using a combination of helicase and 3’ to 5’ nuclease activity digests the DNA until a Crossover hot-spot instigator or Chi sequence (5’GCTGGTGG3’) is encountered. At this point, the RecBCD complex pauses and reverses the polarity of its nuclease activity and creates a 3’ ssDNA over-hand. RecA is then recruited and loaded on the 3’ over-hang by RecBCD complex to initiate strand invasion of the homologous dsDNA[6,19,20]. It is not well understood that in the absence of both RecB and RecC how RecD2 functions in *Deinococcus radiodurans*.

RecD2 from *Deinococcus radiodurans* (DrRecD2) has important contributions towards the organism’s unusually high tolerance to gamma radiation and hydrogen peroxide. *D. radiodurans* can withstand and recover hundreds of double-stranded DNA breaks at 15 kGy gamma irradiation, a condition that is lethal to most living beings including humans. The exact molecular mechanisms behind this phenomenon are not well understood to date, available evidence indicates that the radiation resistance may be attributed to the highly condensed chromosome structure and extremely efficient DNA repair enzymes[7]. The less explored DrRecD2 has sequence homology with RecD at the C-terminal domain, but the extended N-terminal domain and a eukaryotic SRC homology domain 3 (SH3) are unique to this particular enzyme[8]. Atomic-level structural studies confirmed that the beta-hairpin structure of the 1B domain of DrRecD2 is responsible for duplex DNA unwinding and complementary biochemical and biophysical studies reported that the enzyme has a low processivity with one base translocation per ATP hydrolysis[8]. In this study, we have probed the exact binding event and processivity of the enzyme at single-molecule resolution using Protein-induced fluorescence enhancement (smPIFE)[9,10] and Forster resonance energy transfer (smFRET)[11]. Based on the real-time observations we probed the ‘first point of contact’ of DrRecD2 on the DNA and proposed two different binding mechanisms (Collisional Docking and Drifting) of the protein on the ssDNA. We also showed that SSB is able to recruit DrRecD2 on the 5’ overhang and during the DNA unwinding process DrRecD2 is able to displace RecA form the ssDNA.

## Results and Discussion

### Quantification of binding and non-binding interactions of DrRecD2 with the 5’ platform

All the previous studies have mentioned that a 10 to 15 nucleotides long ssDNA is required as the binding substrate for DrRecD2 (molecular weight of approximately 76.4 kDa)[12–14], hence we designed a 12 nucleotides long ssDNA sequence (5’TACAGCTACCTA3’) as the binding platform of DrRecD2 (afterwards referred as 5’ platform). But the available crystal structures of the nucleo-protein complex shows that at one given point of time, only 8 nucleotides can interact with the binding pocket[8], hence, it occurred to us that even within the 5’ platform, there should be a sequential preference of binding in the spatio-temporal context. Therefore, we have attempted to find out the first point of contact of the protein on the DNA using smPIFE. Firstly, after analyzing the inherent secondary structure of the 5’ platform both theoretically and experimentally, using RNAfold Web Server (Institute for Theoretical Chemistry, University of Vienna) and smFRET, we concluded that it forms a hairpin structure (Fig. 1a). When we performed the smPIFE experiments with real-time delivery of DrRecD2, with Cy3 labelled at two different positions of the sequence, one at the 5’ terminal end (Terminal PIFE construct) and the other at the ssDNA-dsDNA junction (Junction PIFE construct) (Fig. 1a & Supplementary Fig. 1), we found that there are two distinct types for interactions in the context of the interaction time scale. We classified them as Tapping, which occurs in millisecond time scale (0.14 ± 0.02 ms) and appears to be random in occurrence, hence, we inferred these to be non-binding events where the protein comes close to the nucleotide sequence and then goes away due to random diffusion or Brownian motion (Fig. 1a, 1b & 1e). The occurrence of this kind of behaviour even with the Internal PIFE construct (Supplementary Fig. 1) where there is no possibility of binding of DrRecD2 confirms that this interaction is indeed non-binding in nature. On the other hand, where we observed a fluorescence enhanced state for a significantly longer time scale i.e. for seconds (4.79 ± 2.00 s), we inferred them as true binding or docking events (Fig. 1e). By careful observations of the smPIFE traces of the docking events, we further classified the interactions into different categories. The binding of DrRecD2 to the DNA can be either through a collision (Collision Docking), where the protein comes to the binding spot very fast by random diffusion and if the DNA binding domain of the protein is rightly oriented towards the DNA at that moment, a fast and stable binding is achieved. These types of events are identified by a very sharp and sudden enhancement of the fluorescence intensity which persists until the probe has photo-bleached or the protein has fallen off the DNA (Fig. 1a & 1c). Another means of binding of the protein on the DNA is by Drifting, where the protein approaches the DNA binding sequence by a #controlled’ slow diffusion process instead of fast collision. In this case, the protein has an ample amount of time to orient its DNA binding domain properly to achieve a successful docking. The typical signature of these types of events is a gradual increase in the fluorescence intensity over time until it reaches the maxima (Fig. 1a & 1d). This mechanism seems to be more regulated and reproducible even in a cellular context and this inference is well supported by our experimental observations. For the experiments performed with the Terminal PIFE construct, 71 percent of the molecules showed Drifting whereas 25 percent of the molecules showed Collisional Docking (We could not confidently classify the bindings of the remaining 4 percent of the molecules). Tapping phenomena were observed in approximately 17 to 20 percent of the molecules which did not show any binding events. For the Junction PIFE constructs, the distribution was 33 percent for both Drifting and Collisional Docking (We could not confidently classify the binding events of the remaining 34 percent molecules). In this case, also, 33 percent of the molecules showed Tapping behaviour (out of those who did not show any binding events). In the case of both Drifting and Collision Docking, after the binding has occurred, the DrRecD2 can either stay fixed at the docking site or it can attain a partial bound state with the DNA where it can diffuse back and forth at the vicinity of the docking site, we have further characterised them as Rigid Docking and Flexible Docking respectively (Fig. 1c), however, we did not quantify them as our observational window was limited by the lifetime of the fluorophore (many molecules photo-bleached just after binding or docking events). From the data, we show that the preferable first point of contact for DrRecD2 is the 5’ terminal end and not the ssDNA-dsDNA junction. One additional point to be noted here is that DrRecD2 can resolve the secondary structure of the 5’ platform in order to bind, this observation is in agreement with previous studies[14].

**Figure 1:**
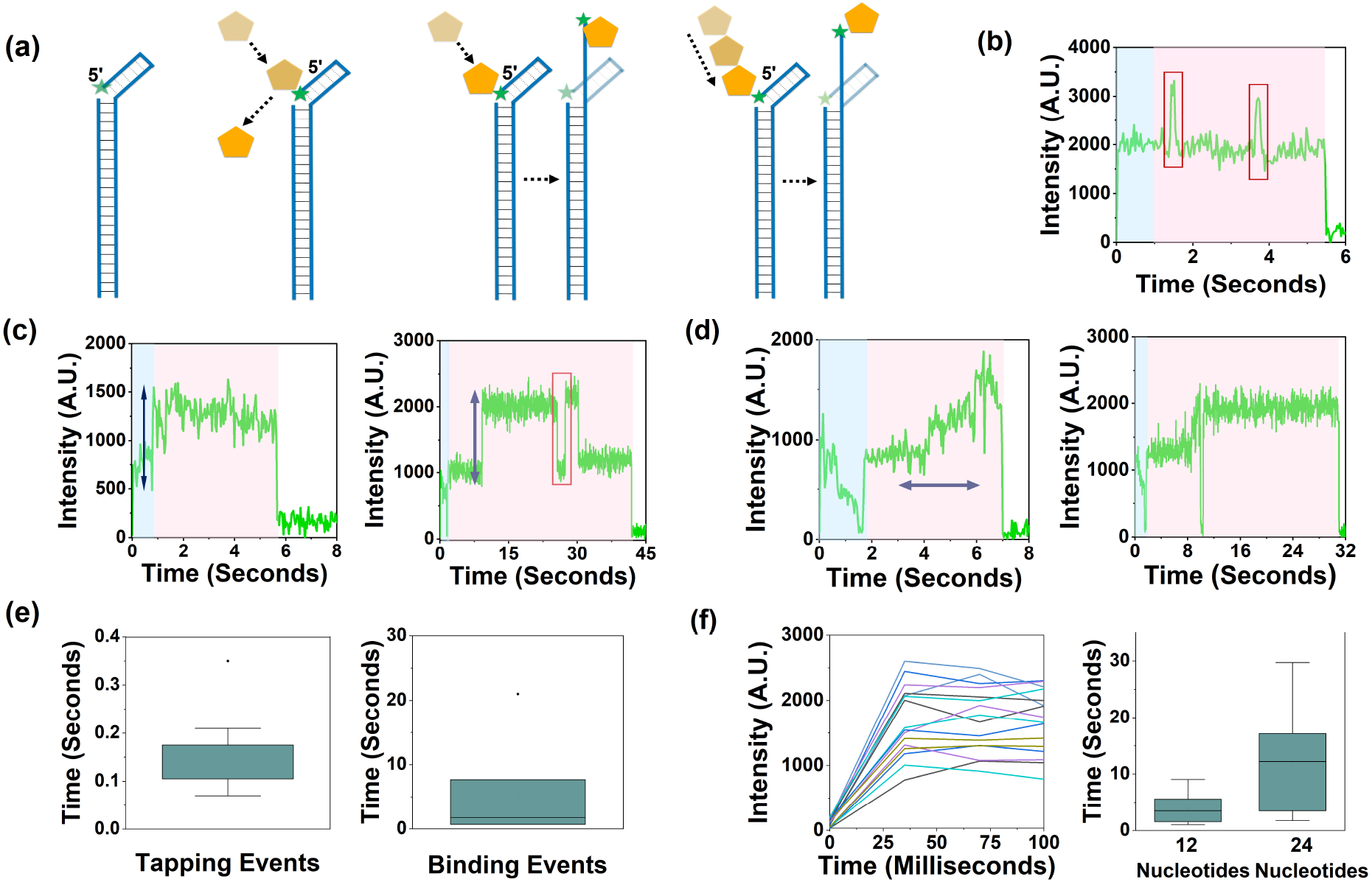
(a) Schematic representation of the different binding mechanisms of DrRecD2 on the 5’ platform. dsDNA and DrRecD2 has been depicted as Blue lines and Yellow Pentagon respectively. Cy3 has been depicted as Green star. The dark Black lines represents hydrogen bonds between the complementary DNA strands and the doted Black lines represents possible electrostatic interactions in the hairpin structure at the 5’ platform. (b) Tapping events are highlighted in Red rectangles. The Blue shaded area represents events before the introduction of DrRecD2 into the sample chamber and the Red shaded area represents the events after the introduction of the protein (this colour code has been followed for rest of the smPIFE traces). (c) Collisional Docking events. The sharp rise in fluorescence has been highlighted in Black arrows. A fall off event has also been recorded and highlighted in Red rectangle. (d) Docking through Drift. The gradual increment of fluorescence has been highlighted with Black arrow. (e) Time scale of Tapping and Binding or Docking events. (f) Time to make the first point of contact with the DNA after the introduction of DrRecD2 into the sample chamber and the translocation time of DrRecD2 to reach 12th and 24th nucleotide.

### Translocation rate of DrRecD2 on ssDNA and dsDNA

To quantify the translocation rate of DrRecD2, we have shifted the Cy3 labelling position 12 nucleotides (Junction PIFE construct) and 24 nucleotides (Internal PIFE construct) away from the 5’ terminal end and have performed smPIFE experiments (Supplementary Fig. 1). As we concluded from our earlier set of experiments that the protein prefers to bind at the 5’ terminal end, hence, any enhanced fluorescence signal away from the 5’ terminal end means the protein has translocated from the terminal towards the fluorophore, and the quantification of the time taken to reach the maximum fluorescence intensity will reflect the translocation rate of DrRecD2. We observed that DrRecD2 takes approximately 35 milliseconds to bind at the 5’ terminal end after introduction into the sample chamber (Fig. 1f). We found a significant difference in the average time scale of reaching the 12th and 24th nucleotide position from the 5’ terminal end, which was 3.88 (± 0.63) seconds and 12.16 (± 3.44) seconds respectively (Fig. 1f). From this data, the physical translocation rate of DrRecD2 on ssDNA and dsDNA was found to be 3.1 nucleotides per second and 1.4 nucleotides per second respectively. Before dsDNA unwinding, the ssDNA enters the binding pocket of DrRecD2 through a narrow cleft formed by its 1B and 2B domains, after this, a series of conformational changes result in additional contacts with domains 1A and 2A which probably allows the beta-hairpin of the domain 1B to locate itself just at the junction of ssDNA-dsDNA[8]. We hypothesise that this elaborate process of loading of DrRecD2 on the ssDNA even before ATP binding might be the reason for the slow rate of translocation on ssDNA. The iterative nature of DNA unwinding in presence of ATP[15] and one-dimensional hopping[16], which are common in many helicases, contributes to the even slower translocation rate on dsDNA.

### Conformational dynamics of the 5’ platform in presence of DrRecD2, SSB, and RecA

We have shown that the 12 nucleotides long 5’ platform attains a hairpin conformation in control experimental conditions i.e. in Reaction Buffer (20 mM Tris-Cl, pH 7.6, 10 mM MgCl2) and from our smPIFE experiments, it was evident that DrRecD2 is capable of resolving this secondary structure to bind. This trend was observed again when we performed smFRET experiments with the Terminal FRET Construct (Fig. 2a & Supplementary Fig. 3). When DrRecD2 was introduced into the sample chamber without ATP or AMP-PNP, the majority of the molecule showed a Low FRET state (75.27 percent), followed by an Intermediate (13.39 percent) and High FRET state (11.34 percent) (Fig. 2c, 2d & Supplementary Fig. 4a). The High and Low FRET indicate protein unbound and bound states respectively, and the observed temporal transition rates from Low to Intermediate (k_LI_=3.2 s^−1^) indicate that the Intermediate FRET states represent the partially folded hairpin structures of the 5’ platform due to the inherent basal level helicase activity of DrRecD2 (Fig. 2e). By combining the results of both smPIFE and smFRET experiments, we conclude that the sequential set of events comprise of (i) binding of RecD2 at 5’ terminal end (ii) sliding along the ssDNA with high speed till ssDNA-dsDNA junction with simultaneous resolution of the secondary structure present and (iii) opening the double stranded DNA with concomitant translocation with a comparative lower speed.

**Figure 2:**
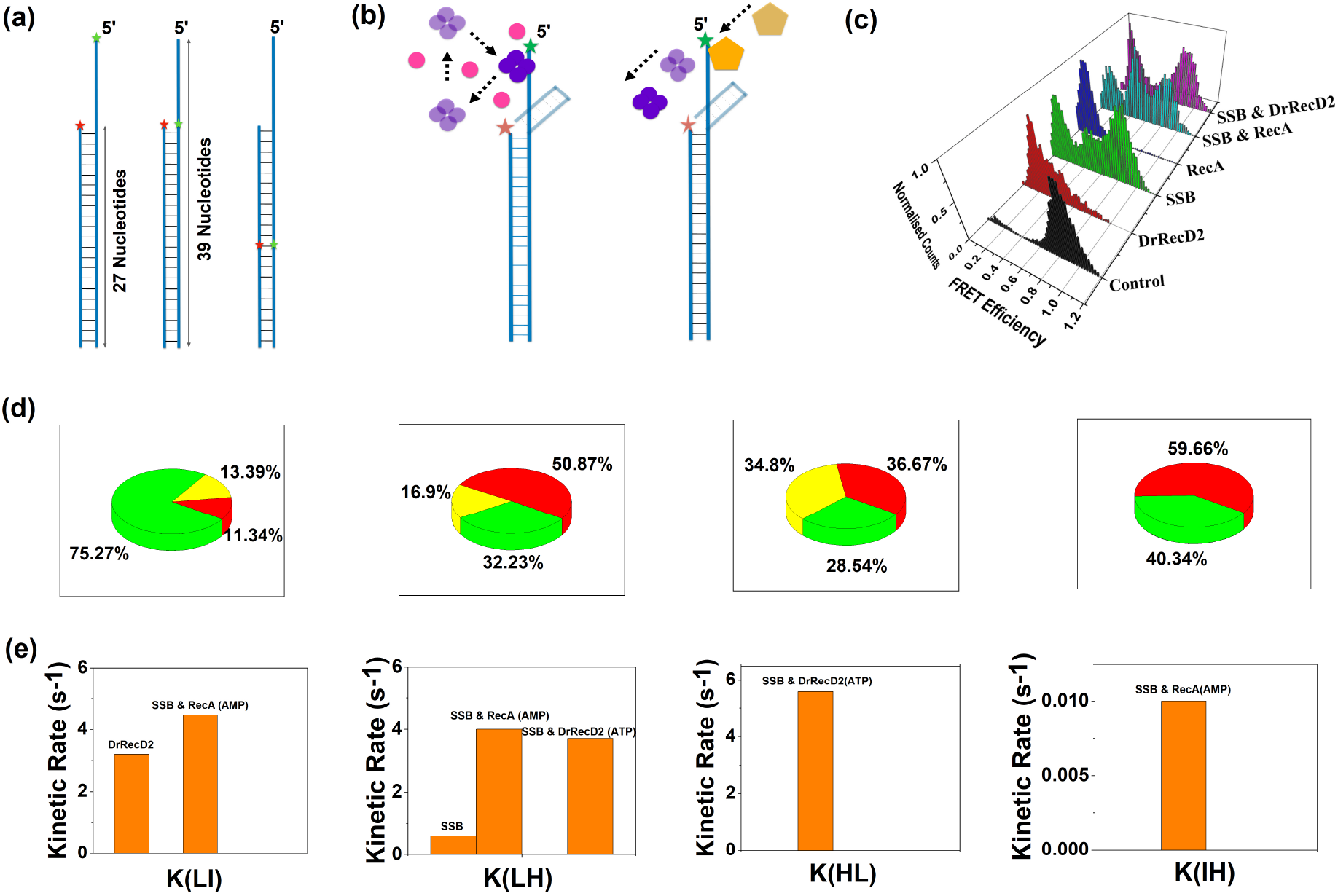
(a) Schematic representation of the Terminal FRET, Junction FRET and Internal FRET Constructs respectively. Same colour codings were followed as before. Cy5 has been represented as Red star. (b) Schematic depiction of competitive binding between SSB & RecA and cooperative binding between SSB & DrRecD2 with the 5’ platform. SSB-homotetramer has been represented as 4 connected Purple circles, RecA has been represented as Pink circles (c) Comparative 3D histograms of FRET efficiencies due to different interactions of proteins with the 5’ platform (d) Pie charts of the population distributions. (e) Bar plots of the kinetic rates.

SSB is known to bind to ssDNA sequences of length 17 to 70 nucleotides as homotetramers (molecular weight of approximately 75.5 kDa) and the binding mechanism is very similar to that of histone proteins in eukaryotes[17,39]. For a 35 nucleotides long ssDNA substrate there is a complete wrapping of the DNA around the SSB-homotetramer, and when the substrate length is either 65 nucleotides or 17 nucleotides there is double wrapping and incomplete wrapping respectively. When SSB was introduced to the 5’ platform, we observed three distinct populations of High (50.87 percent), Intermediate (16.9 percent), and Low (32.23 percent) FRET state (Fig. 2c, 2d & Supplementary Fig. 4c). Similar to our previous observations, the High and Low FRET state denotes the ssDNA unbound and bound the state of SSB-homotetramers, while in this case, the Intermediate FRET state indicates a partially bound state. The binding of SSB-homotetramer on the 5’ platform was independently verified by smPIFE (Supplementary Fig. 2). The transition from Low to High FRET (k_LH_=0.6 s^−1^) state indicates single-step unbinding events of SSB-homotetramers due to short ssDNA substrate (Fig. 2e). SSBs were reported to bind as short as 8 nucleotides long ssDNA by density-gradient centrifugation[40] but previous single-molecule studies did not observe any binding to ssDNA shorter than 17 nucleotides[39]. Hence, this is the first experimental evidence of binding of SSB-homotetramer to short an ssDNA (12 nucleotides) strand at a high salt concentration (>2 mM Mg^+^) at single-molecule resolution.

By contrast to this, when RecA, another ssDNA binding protein was introduced to the same DNA substrate (supplemented with 5 mM AMP-PNP), a single population of Low FRET state was observed (Fig. 2c & Supplementary Fig. 4b). This validates the difference in mechanism of binding of SSB and RecA, while SSB-homotetramer wraps the ssDNA around itself, RecA (molecular weight of approximately 37.9 kDa) binds as a monomer on the ssDNA giving rise to a nucleo-protein filament[18] and as the monomeric units of RecA are smaller compared to the SSB-homotetramers, the substrate length is not a limiting factor for RecA and as a result, a stable binding is achieved. The binding of RecA on the ssDNA was independently verified by smPIFE (Supplementary Fig. 2). Moreover, the length of ssDNA (12 nucleotides) used in this experiment can accommodate only one monomer of DrRecD2 or one unit of the SSB-homotetramer at a time, hence, partial binding by these two proteins can give rise to an Intermediate FRET state which is evident from our experiments. But as one monomer of RecA covers three nucleotides, in principle, four monomers of RecA can bind to the 5’ platform. Probability wise it is highly unlikely for all the four molecules to fall off or stay in a partially bound state at the same time i.e. some RecA molecules are always bound with the ssDNA at any given point of time and hence we do not get any Intermediate or Low FRET state (Fig. 5f).

### Competitive binding of SSB-homotetramer and RecA on 5’ platform

Although RecA is known to displace SSB from long ssDNA[18], the mechanism of this process on short ssDNA has not been explored. Our next objective was to check the nature (competitive vs cooperative) of binding between SSB and RecA on the 5’ platform within the same temporal window. In the cellular context, SSB comes and binds to any available ssDNA to protect it from degradation, and proteins like DrRecD2 and RecA come and displace the SSB-homotetramers at a later stage in the DNA repair pathways[6]. To mimic this condition, we introduced RecA (supplemented with 5 mM AMP-PNP) to SSB-homotetramers bound 5’ ssDNA platform (Terminal FRET Construct) by sequential injection into the sample chamber. RecA was supplemented with AMP-PNP to ensure strong nucleo-protein filament formation. The distribution of the FRET states appeared to be almost uniform across the High (36.67 percent), Intermediate (34.8 percent), and Low (28.54 percent) (Fig. 2d & 2c) but very fast transitions were observed from both Low to Intermediate (k_LI_=4.7 s^−1^) and High (k_LH_=4.0 s^−1^) and slow transitions from Intermediate to Low (K_IL_=0.01 s^−1^) were also observed (Fig. 2e & Supplementary Fig. 5a). These observations are in complete contrast with our previous SSB-homotetramer bound conditions, the fast transitions from Low to Intermediate and High indicate RecA can displace the SSB-homotetramer from the 5’ platform either partially or completely. The slow transitions from Intermediate to High in some of the molecules suggest that the complete displacement of SSB-homotetramer by RecA preferably occurs in a single step but in some cases, it can happen through an intermediated state (two-step). Surprisingly we did not observe any reverse transitions and the distribution of the FRET states remained unaltered throughout the observational window of our experiments. This led us to conclude that displacement of SSB-homotetramer from ssDNA by RecA and binding of RecA on the same substrate are mutually independent. From our data, it is evident that these two competitive events are not well coordinated and there is indeed a significant delay in RecA binding on the ssDNA after SSB-homotetramer displacement (Fig. 2b). This observation supports the hypothesis that *in-vivo*, RecA is assisted by RecFOR and RecOR to displace SSB-homotetramer from the ssDNA and bind on it[27–29].

### Cooperative binding of SSB-homotetramer and DrRecD2 on 5’ platform

Keeping the same objective and logic of competitive binding and displacement in mind we checked the behaviour of SSB and DrRecD on the 5’ platform (Terminal FRET Construct). When DrRecD2 was introduced to the SSB-homotetramer bound 5’ platform, distinct High (59.60 percent) and Low FRET (40.34 percent) states were observed (Fig. 2d & Supplementary Fig. 5b), which are absolute contrast with our previous experiments with only DrRecD2 and SSB, wherein both the cases we got a significant amount of Intermediates FRET states (Fig. 2c). Not only that, in this case, for the first time we observed transitions both from High to Low (k_HL_=5.8 s^−1^) and reverse (k_LH_=3.7 s^−1^) (Fig. 2e). The Low FRET state represents the bound state of either SSB-homotetramer or DrRecD2 on the 5’ platform and the transition for Low to High FRET state indicates that DrRecD2 is displacing the SSB-homotetramer and making the 5’ platform free to fold back into its native hairpin structure. We rule out the possibility of self dissociation of SSB-homotetramer (k_LH_=0.6 s^−1^) based on the observed fast temporal transitions (k_LH_=3.7 s^−1^) (Fig. 2e). We also rule out the possibility of transition from Low to High FRET state by the basal level helicase activity of DrRecD2 because as evident from our previous data, it happens through an Intermediate FRET state. The follow-up rapid reverse transition from High to Low FRET state (k_HL_=5.8 s^−1^) indicates binding of DrRecD2 (Fig. 2e). From the observations, we conclude that unlike poorly coordinated SSB-RecA competitive binding, the SSB-DrRecD2 binding events are mutually dependent, well-coordinated, and cooperative in nature (Fig. 2b). Although no direct experimental evidence of interaction of SBB and DrRecD2 are found to date, the interactions of other helicases from the Rec family-like RecG, RecQ, RecJ, and RecO with SSB are well established[24–26]. The rapid association of DrRecD2 on the 5’ platform in presence of SSB favours the possibility of recruitment of DrRecD2 by the SSB-homotetramer in actual *in-vivo* conditions.

### Effect of DrRecD2 at the ssDNA-dsDNA junction

In the context of double-stranded DNA break repair, ssDNA-dsDNA junction regions are always the starting point of the helicase activity of DrRecD2. Hence we checked the immediate downstream effects of binding of DrRecD2 at the 5’ platform by putting the FRET pair (Cy3 and Cy5) exactly opposite to each other at the ssDNA-dsDNA junction (Junction FRET Construct) (Fig. 2a). To our surprise, we did not find any High FRET state as one would expect when the control experiments were performed with Reaction Buffer (20 mM Tris-Cl, pH 7.6, 10 mM MgCl2), a stepwise transition from a High to Low FRET state was also observed in some molecules. However, a significant amount of the High FRET population appeared when the Reaction Buffer was supplemented with 5 mM ATP, suggesting that the junction is getting stabilised (Supplementary Fig. 6). A thorough literature survey gave us two possible explanations of this behaviour, firstly, complementary nucleotides at the ssDNA-dsDNA junctions can undergo partial unwinding, a phenomenon known as molecular fraying, where there is transient destabilisation of the terminal base pairs of DNA or RNA due to thermal fluctuations[38.] Secondly, when two fluorophores are present in very close proximity (<5 nm) with each other (similar to our condition) there is a high chance of getting low fluorescence signal due to collisional quenching or self-aggregation of the fluorophores with each other[21–23].

When DrRecD2 was introduced into the sample chamber without any ATP or AMP-PNP, a significant population in the Low FRET (60.32 percent) state was observed which was due to the inherent basal level helicase activity of DrRecD2. A range of FRET states was observed from 0.5 to 1.0 which can be further segregated into Intermediate (8.4 percent), Intermediate-High (11.08 percent), and High (20.2 percent) upon careful inspection (Fig. 3a, 3b & Supplementary Fig. 7a). Molecules showing Intermediate and Intermediate-High FRET states represent the conditions where DrRecD2 is showing its helicase activity at the ssDNA-dsDNA junction and after that, it is translocating away from the junction, once the protein is far away from the junction the dsDNA can rewind, which is getting reflected as High FRET state. In the same experiment when performed in presence of 5 mM AMP-PNP, distinct High (77.98 percent) and Low (22.02 percent) FRET states were observed. Only Low FRET (100 percent) state was observed when supplemented with 5 mM ATP (Fig. 3a & 3b and Supplementary Fig. 7b & 7c). The disappearance of Intermediate FRET states and fast temporal transition from Low to High FERT state in presence of 5 mM AMP-PNP compared to only DrRecD2 condition (k_LH_=5.4 s^−1^ and 0.8 s^−1^ respectively) indicates that the protein can move away from the junction faster in presence of AMP-PNP but unable to unwind the dsDNA completely (Fig. 3c). The fact that we got only Low FRET state in presence of ATP indicates that DrRecD2 is able to unwind the dsDNA completely and as a result, the Donor-fluorophore (Cy3) labelled strand got completely dissociated from its complementary Acceptor-fluorophore (Cy5) labelled strand. The Low FRET state (38.3 percent) reappeared when DrRecD2 was supplemented with 5 mM ATP and SSB (Fig. 3a, 3b & Supplementary Fig. 7d). The high temporal transition from High to Low FRET (k_HL_=12.8 s^−1^) state in this condition again suggests the possible recruitment of DrRecD2 by SSB (Fig. 3c) and the association of SSB with the freshly unwinded ssDNA should slow down the reverse transition (i.e. from Low to High) in principle, and this is what exactly getting reflected in our data (k_LH_=0.2 s^−1^) (Fig. 3c). Thus, our observations indicate that, in presence of ATP, DrRecD2 is highly processive, separating both the strands of the DNA, however even without ATP there is unwinding as well as translocation of the protein along the dsDNA track.

**Figure 3:**
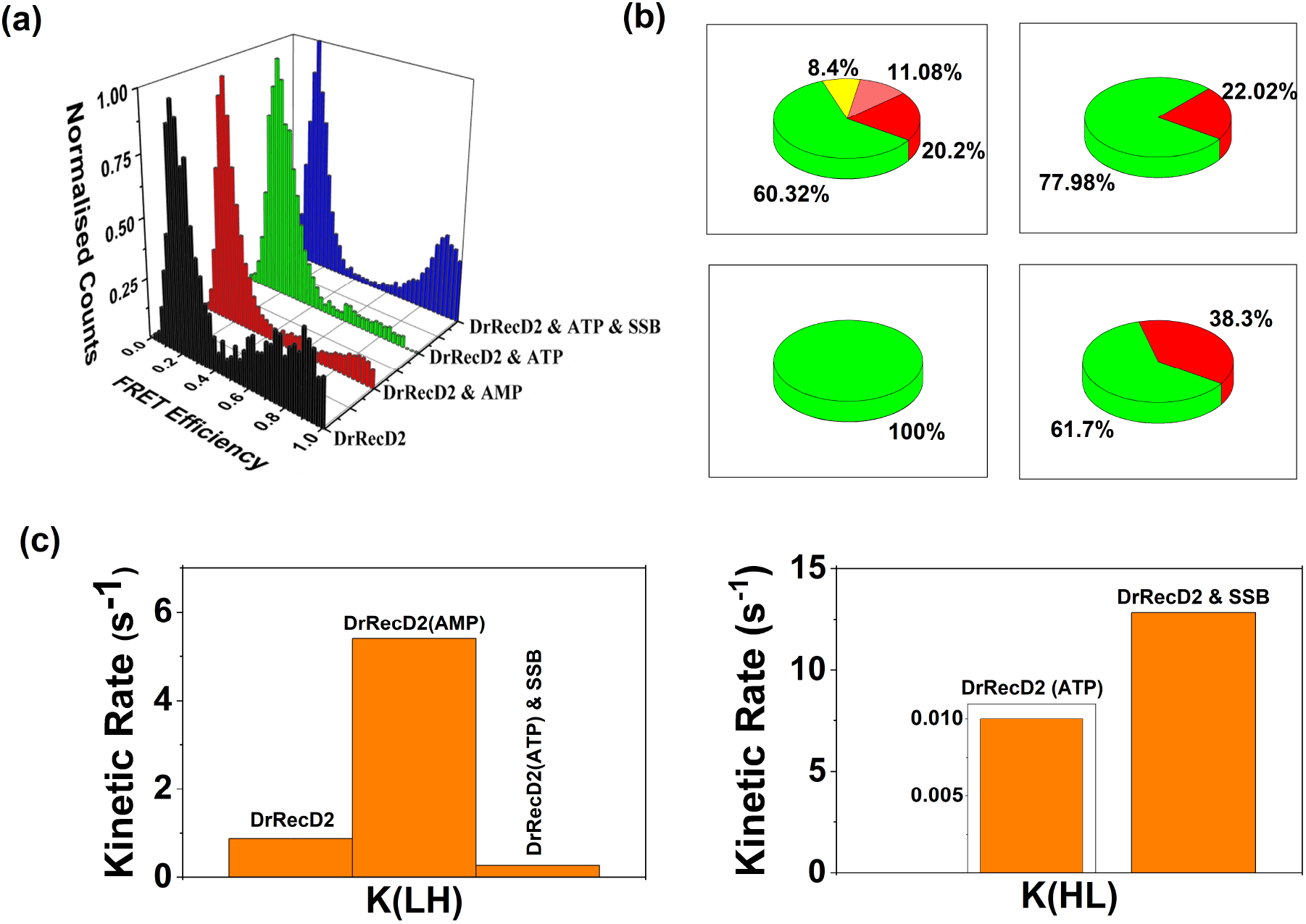
(a) Comparative 3D histograms of FRET efficiencies due to different interactions of proteins with the Junction FRET Construct. (b) Pie charts of the population distributions. (c) Bar plots of the kinetic rates.

### The heterogeneous unwinding of dsDNA by DrRecD2

To monitor the dsDNA unwinding ability of DrRecD2 unambiguously, we placed the FRET pair opposite to each other 24 nucleotides away from the 5’ terminal end (Internal FRET Construct) (Fig. 2a & Supplementary Fig. 8). In presence of only DrRecD2, a large population in the Low FRET (67.51 percent) state along with Intermediate (22.49 percent) and High (10 percent) was observed (Fig. 4a, 5a & 5b). Fast transition from High to Intermediate FRET state (k_HI_=3.1 s^−1^) followed by a slower reverse transition (k_IH_=0.7 s^−1^) and absence of any transition from Low to either Intermediate or High FRET indicates differential basal level helicase activity within different molecules of DrRecD2 (Fig. 4d & 4c). Even without ATP some of the molecules are able to completely unwind the dsDNA, completely displacing the Donor-fluorophore (Cy3) labelled strand out in the solution, while the rest of the molecules are displaying partial or incomplete unwinding (Fig. 5e). To our knowledge, this heterogeneous behaviour of DrRecD2 even in absence of ATP has not been reported earlier. The population distribution of the FRET efficiencies changes drastically (52.91 percent High, 14.68 percent Intermediate, and 32.41 percent Low) when DrRecD2 is supplemented with 5 mM AMP-PNP (Fig. 4a, 5a & 5b). The increment in the High FRET state with rapid forward (k_HI_=16 s^−1^) and reverse (k_IH_=14.6 s^−1^) transition from High to Intermediate FRET state indicates iterative nature of DNA unwinding (Fig. 4d & 4c). Transitions directly towards the Low FRET state from the High FRET (k_HL_=0.4 s^−1^) state by some of the molecules reflects functional heterogeneity in the system (Fig. 4d). The signatures of the iterative nature of DNA unwinding were captured in detail when DrRecD2 was supplemented with 5 mM ATP. The population in the High FRET state decreased (24.74 percent), followed by an increased Intermediate (31.85 percent) and Low (43.41 percent) FRET state population (Fig. 4a, 5a & 5b). Fast transition from High to Intermediate FRET state (k_HI_=4 s^−1^) followed by two transitions from Intermediate FRET state to either Low FRET state (K_IL_=0.3 s^−1^) or back to High FRET state (k_IH_=2.5 s^−1^) indicate that most of the DrRecD2 molecules are falling off from the DNA strand before reaching the 24th base-pair while unwinding it (Fig. 4d & 4c) and the transition from Low to Intermediate FRET state (k_LI_=0.7 s^−1^) (Fig. 4c) indicates the few molecules those reach at the 24th base-pair (or beyond that) before falling off, are unable to unwind the DNA completely, otherwise, we would have seen no back transition from Low to either Intermediate or High FRET state (Fig. 5e).

**Figure 4:**
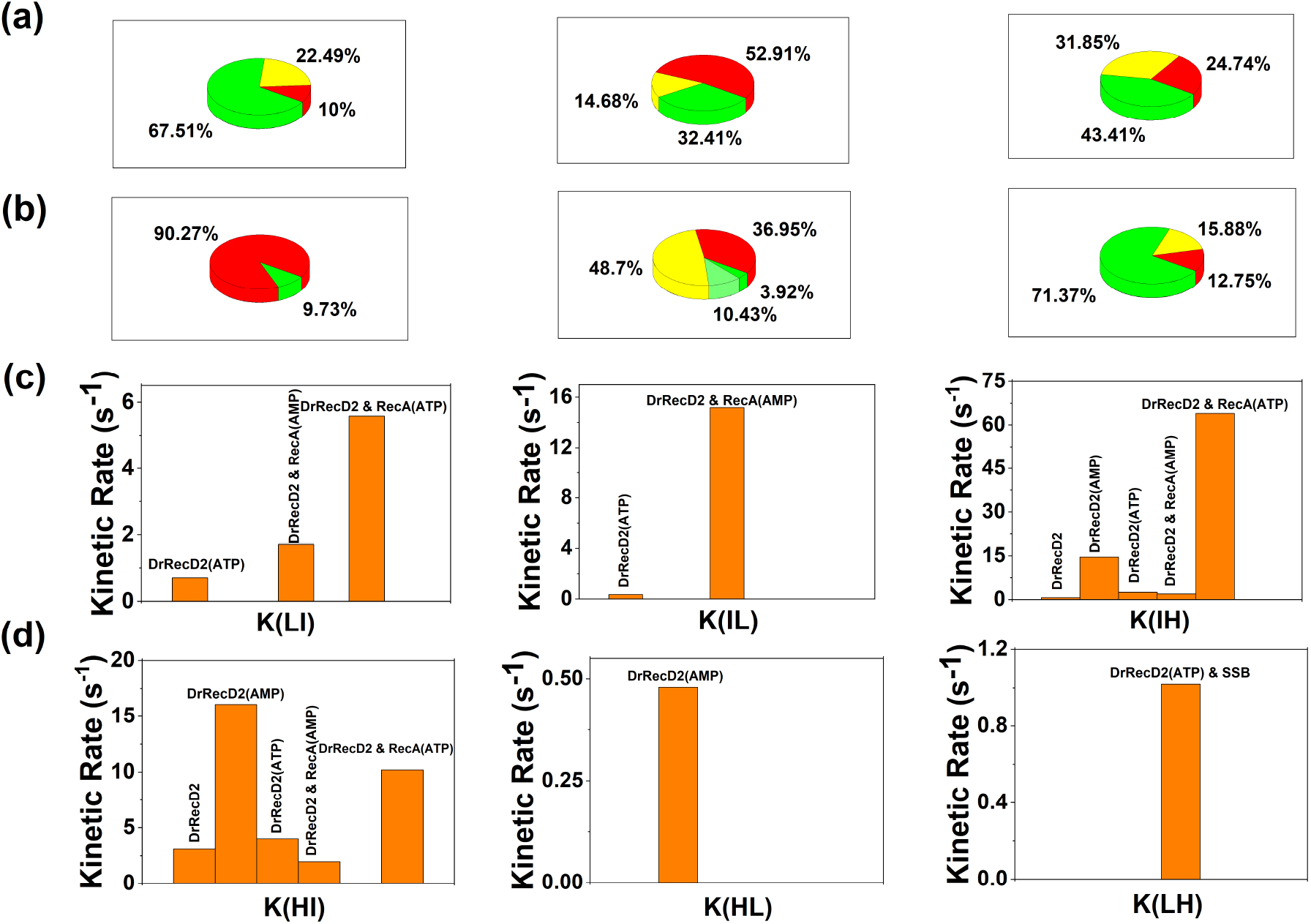
Quantification of FRET efficiency distribution and the kinetic rates for the experiments done with the Junction FRET Construct. (a) & (b) Pie charts of the population distributions. (c) & (d) Bar plots for the kinetic rates.

**Figure 5:**
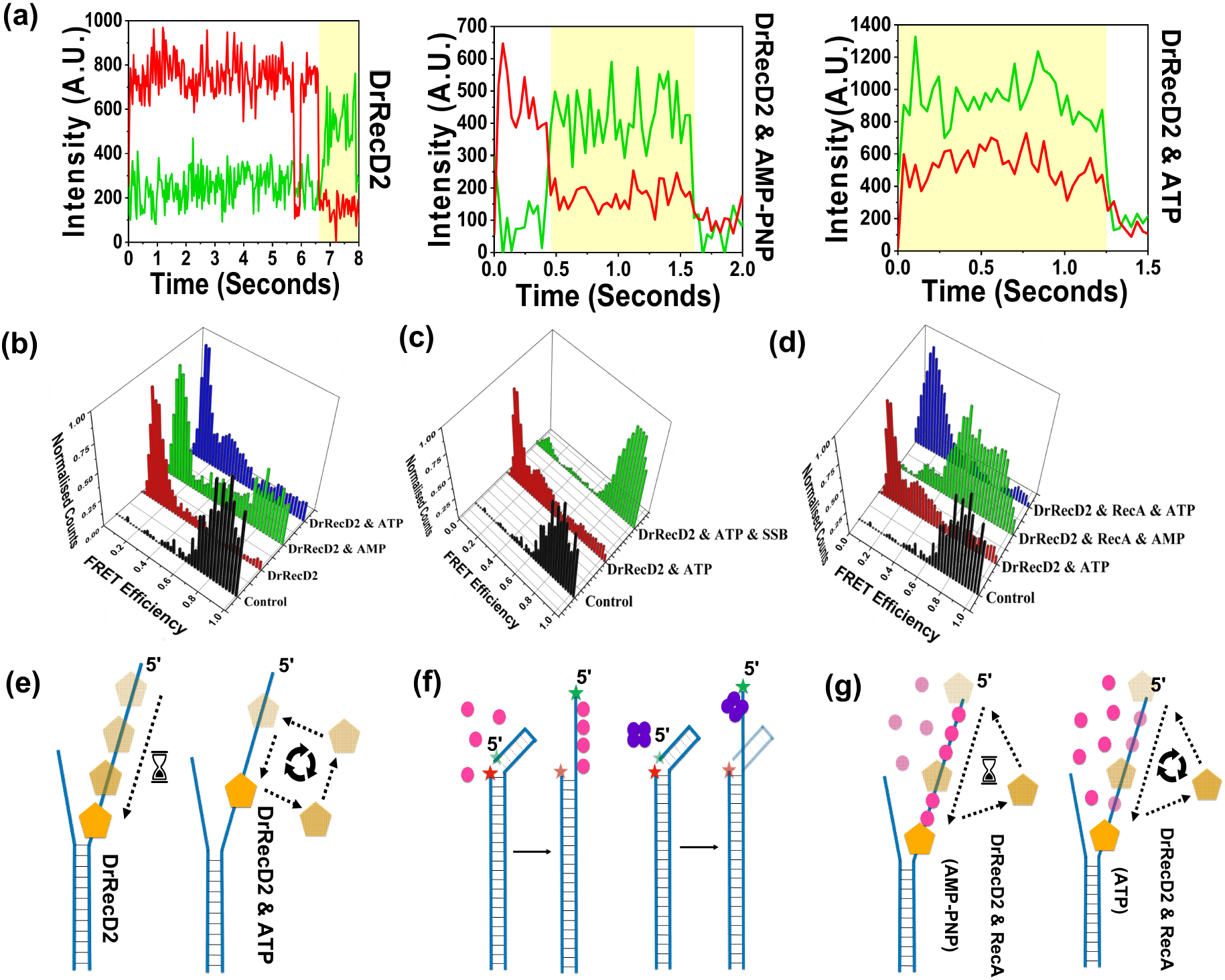
(a) smFRET traces showing the initiation point of dsDNA unwinding by DrRecD2 in temporal context. The High FRET state indicates stable dsDNA and the Low FRET state highlighted in Yellow shade indicates the unwinding event by the protein. (b-d) Comparative 3D histogram of FRET efficiencies resulting from differential protein activities on the Internal FRET Construct. (e) Schematic representation of DNA unwinding by DrRecD2. (f) Schematic representation of difference in binding modes of RecA and SSB. (g) Schematic representation of interaction of DrRecD2 with RecA filaments.

### Partial DNA unwinding by DrRecD2 in presence of SSB

The unwinding of dsDNA by DrRecD2 produces longer ssDNA which can act as a substrate for the SSB-homotetramer, in the process if DrRecD2 dissociates from the DNA then it may not be able to bind again as the DNA will be pre-occupied by the SSB-homotetramer. To check this hypothesis, we introduced both DrRecD2 and SSB simultaneously, supplemented with 5 mM ATP into the sample chamber containing the immobilised DNA (Internal FRET Construct). We observed distinct High (90.27 percent) and Low (9.73 percent) FRET states with molecules transiting from Low to High (k_LH_=1 s^−1^) FRET states (Fig. 4b, 5c & 4d). The absence of transition from High to Low (k_HL_) FRET, again suggests rapid DrRecD2 recruitment by SSB-homotetramer followed by DNA unwinding. The DNA molecules where DrRecD2 got dissociated from the DNA due to its iterative nature of unwinding showed High FRET as the binding site of the helicase i.e. the 5’ platform is now completely occupied by the SSB-homotetramer and hence DrRecD2 is unable to bind, leaving the dsDNA partially unwinded. From these experiments we conclude that once DrRecD2 has unwinded the dsDNA till 24^th^ nucleotide, the entire 24 nucleotide long single-stranded sequence is now available for SSB-homotetramer to bind. Therefore, once DrRecD2 has fallen off the DNA while unwinding, it is unable to displace the SSB-homotetramer from the ssDNA.

### Displacement of RecA from the DNA by DrRecD2

Unlike the histone or SSB, RecA is known to form strong DNA filaments in presence of AMP-PNP, whereas in presence of ATP the filaments are found to be weak or transient in nature as the RecA monomers tend to dissociate from the ssDNA after each ATP-hydrolysis event[30–32]. We wanted to check if DrRecD2 can disrupt the RecA-mediated DNA filaments *in-vitro*. RecA, supplemented with 5 mM AMP-PNP was introduced into the sample chamber containing the immobilised DNA (Internal FRET Construct) and was incubated for 5 minutes to ensure strong filament formation after that DrRecD2 was injected into the sample chamber. Approximately one-third of the molecules (36.95 percent) showed High FRET, indicating DrRecD2 is unable to bind to the 5’ platform. However, the presence of Intermediate (48.7 percent), Low (10.43 percent), and Very-Low (3.92 percent) FRET states does indicate DNA unwinding by DrRecD2 after displacing the RecA monomers from the 5’ platform (Fig. 4b, 5d & Supplementary Fig. 9a). The forward and reverse transitions, both from High to Intermediate (k_HI_=1.9 s^−1^ and k_IH_=2 s^−1^) and from Intermediate to Low (K_IL_=15.2 s^−1^ and k_LI_=1.7 s^−1^) reconfirms the iterative nature of helicase activity of DrRecD2 (Fig. 4d & 4c). The clear distinction among the molecular populations at the low FRET region suggests that the process of DNA unwinding in presence of strong RecA filaments is not very smooth and happens in a stepwise manner. It should also be noted that DrRecD2 is unable to utilise its full potential as we have used AMP-PNP instead of the natural substrate ATP (Fig. 5g).

When the same experiment was performed supplemented with 5 mM ATP, a drastic change in FRET distribution was observed. Almost three-fold decrement in the High FRET state (12.75 percent) and five-fold increment in the Low FRET state (71.37 percent) was observed which clearly indicates an efficient DNA unwinding by DrRecD2 (Fig. 4b, 5d & Supplementary Fig. 9b). Reduction in Intermediate FRET state (15.88 percent) and rapid transitions from High to Intermediate FRET state (k_HI_=10.2 s^−1^) and reverse (k_IH_=64 s^−1^) (Fig. 4d & 4c) demonstrate efficient iterative displacement of RecA from the weak filament by ATP-coupled DrRecD2 while unwinding the DNA (Fig. 5g).

## Conclusions

From our experiments, we showed that DrRecD2 prefers to approach the 5’ terminal end in a slow and controlled manner to bind. The protein has the ability to resolve secondary structures like hairpins and its recruitment is enhanced in presence of SSB. The rate of translocation of DrRecD2 appears to be slower on the dsDNA compared to ssDNA, which is due to the iterative nature of DNA unwinding by DrRecD2 in presence of ATP. We showed that DrRecD2 is able to unwind dsDNA even in absence of ATP or AMP-PNP at a slower pace, however, the iterative mode of DNA unwinding has also been reported in Rep helicases[33], is evident only in presence of ATP or AMP-PNP. The inherent helicase activity of DrRecD2 without ATP suggests that on a naked dsDNA, DrRecD2 is able to carry out its function smoothly, however, in the actual cellular context where the DNA is always in association with other proteins (i.e. in chromatin state), the ATP-hydrolysis becomes essential to unwind the duplex while simultaneously displacing the other proteins. It is also possible that the energy of ATP-hydrolysis by DrRecD2 is used by a multi-protein assembly complex as has been reported in the case of DnaB helicase[34]. By looking at the interaction of DrRecD2 with RecA-mediated DNA filaments, it occurred to us that the energy released by the ATP-hydrolysis from DrRecD2 can also be channelised to regulate the RecA filament dynamics. Similar observations have been reported where the energy released by ATP-hydrolysis by RecN is altering the dynamics of RecA-mediated DNA filaments[35]. The function of ATP-hydrolysis by RecA is still a mystery to the scientific community as RecA is neither a helicase nor a translocase protein. The nature of the interaction between RecA and DrRecD2 observed in our experiments may shed some light on why RecA evolved having an ATP binding domain. In order to make space for other proteins to bind during the DNA repair process, the RecA-mediated DNA filaments have to be transient. Hence, we hypothesise that the micro-organisms having a RecA variant without the ATP binding domain were inefficient in repairing their damaged DNA as the strong RecA-mediated DNA filaments did not allow much room for other essential repair proteins to come and bind and a result they eventually faced extinction. On the other hand, some mutants having RecA with an ATP binding domain happened to counter any dsDNA breaks more efficiently due to their transient RecA-mediated DNA filaments as this allowed other accessory proteins to bind to the DNA and carry out their respective functions. On a similar note, during the RecA mediated homology search, the nature of the filament has to transient in order to facilitate the hydrogen binding between the complementary homologous DNA strands.

We also demonstrated that DrRecD2 is unable to displace the SSB-homotetramer when it is tightly associated with a longer ssDNA substrate and as a result, the dsDNA remains partially unwinded. On the other hand, RecA can be displaced by DrRecD2 even from the strong DNA filaments. These observations open up the intriguing possibility of a coalition of SSB-RecA-DrRecD2 during the homologous recombination of double-strand DNA break repair pathways. The facilitated loading of RecA at Chi sequence by RecBCD complex has already been reported[36]. We hypothesise that when a double-strand DNA break occurs in a cell, at the initial stage when the ssDNA is short, DrRecD2 is recruited at the site by the SSB-homotetramers. Over time as the helicase unwinds the dsDNA, the SSB-homotetramers bind to the newly exposed ssDNA and protect it from degradation. At this stage, RecA comes and starts binding on the ssDNA after displacing the SSB-homotetramers. In this situation even if DrRecD2 falls off from the DNA, it is able to bind to it again by displacing the RecA monomers, as a result, slow and iterative unwinding of the DNA is achieved along with the formation of transient RecA-mediated DNA filaments, which eventually leads to homologous strand searching followed by recombination (Fig. 6).

**Figure 6:**
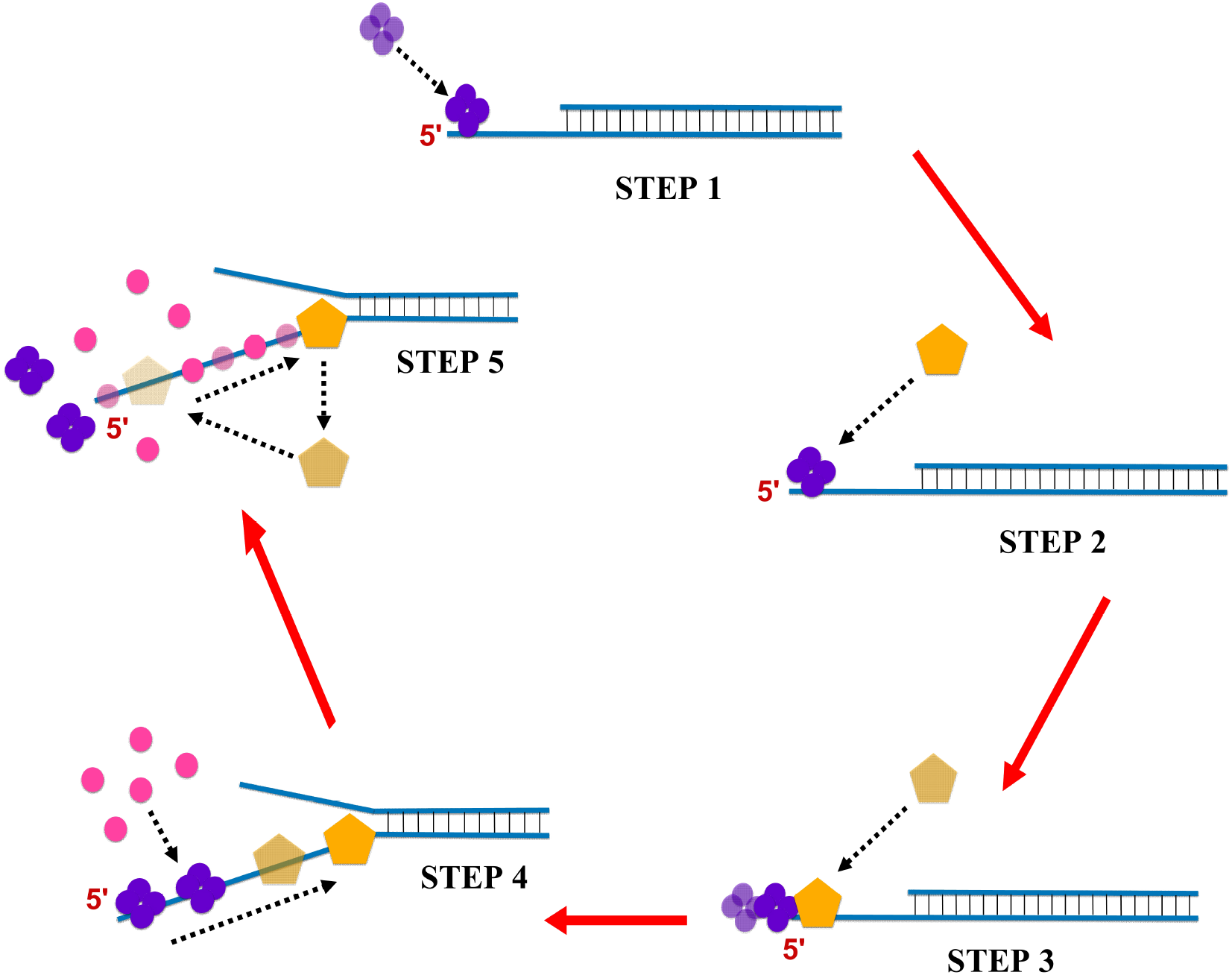
The final schematic model of the possible sequential interaction of SSB, DrRecD2 and RecA in DNA repair pathway.

## Materials and methods

All the reagents used for single-molecule experiments were of analytical grade and were purchased from Sigma-Aldrich and Merck. The common salts used in protein purification were purchased from Sisco Research Laboratories Pvt. Ltd. (SRL) and media for bacterial cell culture were purchased from HiMedia Laboratories. All the buffers and media were made with ultra-purified water from Arium Comfort Series, Sartorius.

### Induced over-expression DrRecD2

RecD2 gene from *Deinococcus radiodurans* cloned in plasmid having Kanamycin antibiotic marker and a 6 Histidine tag (His-tag), was a generous gift from Prof. Juan C. Alonso, Department of Microbial Biotechnology, CNB-CSIC[41]. The plasmids were transformed into *Escherichia coli* BL21 (DE3) cells and kept for 12 hours at 37°C after spreading it on Luria-Bertani agar plates supplemented with 50 ug/ml Kanamycin. For protein expression, fresh colonies transformed into *Escherichia coli* BL21 (DE3) cells were inoculated in 1 L of Luria-Bertani broth media supplemented with 50 ug/ml Kanamycin and the culture was grown at 37°C for 2.5 to 3 hours until the absorbance of the culture reached at 0.4 at 600 nm. At this point, induction was done with 2 mM of Isopropyl ß-D-1-thiogalactopyranoside (IPTG), after that the culture was grown for another 3 hours at 37°C. Then the culture media was centrifuged (Heraeus, Biofuge, Stratos, Thermo Fisher Scientific) at 4000 rpm at 4°C for 15 min and the cells were collected. The cells were washed with ultra-purified water supplemented with 10% Glycerol repeatedly three times with centrifugation keeping all the parameters the same and subsequently stored at −20°C.

### Purification of DrRecD2

The protein of our interest, DrRecD2 having a molecular weight of approx. 76.4 kilo-Dalton (kDa) was purified with high purity using Immobilized Metal Affinity Chromatography (IMAC). For lysis, cells were first resuspended in Lysis Buffer (50 mM Tris-Cl, pH 7.6, 500 mM NaCl, 2 mM PMSF, 10% Glycerol, 0.4 mg/ml Lysozyme) and then sonicated (Ultrasonic Processor, Cole Parmer) with 5 cycles of 30 s #ON’ and 30 s #OFF’ sonic pulses keeping 14% amplitude. The cell lysate was centrifuged at 10,000 rpm at 4°C for 1 hour, the collected supernatant was manually loaded into a pre-equilibrated (with Lysis Buffer) Ni-NTA affinity chromatography column and allowed to pass through it with a flow rate of 0.5 ml/min (this flow rate was maintained throughout the entire experiment unless stated otherwise). The column was washed thoroughly with 6 CV (column volume) of Wash Buffer (20 mM Tris-Cl, pH 7.6, 500 mM NaCl, 50 mM Imidazole, 1 mM DTT, 10% Glycerol). The elution was done with 10 ml of Elution Buffer (20 mM Tris-Cl, pH 7.6, 500 mM NaCl, 500 mM Imidazole, 1 mM DTT, 10% Glycerol). The presence and purity of the protein was checked by Bradford Assay followed by Polyacrylamide Gel Electrophoresis (SDS-PAGE, 5% Stacking, and 12% Resolving). After that, the buffer exchange was performed with Dialysis Buffer (20 mM Tris-Cl, pH 7.6, 150 mM NaCl, 1 mM DTT) and simultaneously concentrated using 50 kDa Amicon Ultra-centrifugal Filters (Merck) followed by a final quality check by SDS-PAGE (5% Stacking and 12% Resolving). The samples were then aliquoted into 100 ul fractions in micro-centrifuge tubes and stored at 4°C. The protein was consumed in experiments within 3 to 4 days after purification to avoid aggregation and denaturation. The entire purification progress was performed at 4°C. The working concentration of DrRecD2 was kept at 100 nM in all experimental conditions unless stated otherwise.

### Purification of Single-strand Binding Protein (SSB)

SSB protein from *Escherichia coli* was purchased from Sigma-Aldrich/Merck, Cat. No. S3917. The working concentration of SSB was kept at 50 nM in all experimental conditions unless stated otherwise.

### Purification of Recombinase A (RecA)

RecA protein from *Escherichia coli* was purchased from New England BioLabs (NEB), Cat. No. M0249L. The working concentration of RecA was kept at 300 nM in all experimental conditions unless stated otherwise.

### Design of DNA constructs

Custom-made HPLC purified fluorescent labelled and unlabelled single-stranded oligonucleotides were purchased from Integrated DNA Technologies Inc. (IDT), Coralville, USA. Keeping the original DNA sequence the same, 6 different #working combinations’ of dsDNA having 12 nucleotides over-hand at 5’ end were generated by differential positional labelling of the fluorophores. All the #working combinations’ of dsDNA were generated keeping in mind the experimental conditions for smFRET and smPIFE. Cyanine 3 (Donor) and Cyanine 5 (Acceptor) were chosen as FRET pairs. A short biotinylated strand of 15 nucleotides was also designed for the surface immobilization of the dsDNA. As a final product after annealing, it consisted of a double-stranded region of length 27 nucleotides and a single-stranded region of length 12 nucleotides. The sequences of the fluorescent labelled dsDNA have been described in detail in the supplementary information (Supplementary Table 1).

To carry out the experiments appropriate oligonucleotide strands were selected along with the biotinylated strand. The molar ratios were kept as Biotin:Cy3:Cy5 = 1:1:1 (for smFRET) and Biotin:Cy3:unlabelled strand = 1:1:1 (smPIFE). After mixing the individual single-stranded DNA in T50 Buffer (10 mM Tris-Cl, pH 8.0, 50 mM NaCl), the mixture was heated up to 95°C followed by a controlled cooling down to 4°C (peqSTAR Thermocycler, PEQLAB Biotechnologie) to ensure efficient annealing. The final concentration of the annealed product was kept at 10 uM. During the experiments, the final #single-molecule’ concentration (pico-molar level) was achieved by further dilution of the annealed product in Reaction Buffer (20 mM Tris-Cl, pH 7.6, 10 mM MgCl2). The remaining annealed product was stored at −20°C for future use.

### Single-molecule experiments (smFRET and smPIFE) and data acquisition

Quartz slides were drilled and chemically treated to facilitate surface immobilization of biotinylated samples as explained in detail in our previous study[37]. The details of the data acquisition process and specifications about the instrumental setup are the same as described in our previous paper[37]. Streptavidin (0.2 mg/ml) was introduced into the sample chamber and after 5 min of incubation, the unbound fraction of Streptavidin was washed off by T50 Buffer. Then the biotinylated samples were introduced into the Streptavidin-coated sample chamber, after 5 min of incubation a short pulse of laser (532 nm) was given in order to check the approximate number of immobilized samples, if not satisfied, samples may be reintroduced with higher concentrations. After adjusting the number of immobilized molecules to approximately 400, the unbound samples were washed off by Imaging Buffer (Reaction Buffer supplemented with 1 mM Trolox, 1 ug/ml Glucose, 1 mg/ml Glucose oxidase, and 0.03 mg/ml Catalase). For smFRET experiments, at this point, 100 nM of DrRecD2 and 5 mM of ATP (in Imaging Buffer) was introduced into the sample chamber and data acquisition was initiated. For smPIFE experiments, only 100 nM of DrRecD2 (in Imaging Buffer) was introduced into the sample chamber. smPIFE experiments were performed in both #Incubation Mode’ and #Real-time Mode’. For #Real-time Mode’, DrRecD2 was introduced into the sample chamber after a certain time frame while the data acquisition was still going on, whereas for the #Incubation Mode’, the data acquisition started after the introduction of DrRecD2 into the sample chamber. SSB and RecA were also introduced into the sample chamber as per experimental requirements. All the experiments were performed at room temperature (25°C).

### Data analysis

From the smPIFE data, approximately 30 molecules were chosen form each set of experiments that showed single-step photo-bleaching and a binding event. Each traces were carefully analysed by custom made IDL based codes written in C and the out comes were plotted in Origin 2018.

For the smFRET data, approximately 200 molecules were selected from each set of experiments based on single-step photo-bleaching and donor-acceptor anti-correlation. Each traces were carefully analysed by custom made IDL based codes written in C and the out comes were plotted in Origin 2018. The data analysis pipeline has been mentioned in detail in our previous paper[37].

## Supporting information

DrRecD2_Supplementary_Data

## Acknowledgements

The authors would like to thank Debolina Bandyopadhyay for her critical comments on the manuscript, Manali Basu for her technical help during the data analysis and Soumen Mandal and Saikat Sadhukhan for providing the Origin 2018 software. The authors are grateful for the generous funding support given by the Department of Atomic Energy (DAE) and Council of Scientific and Industrial Research (CSIR), Govt. of India.

## Conflict of Interest

The authors declare that they do not share any conflict of interest.

## Notes

### Competing Interest Statement

The authors have declared no competing interest.

