## Supplementary material for "Probing the molecular mechanism of the action of *Deinococcus radiodurans* RecD2 at single-molecule resolution": DrRecD2_Supplementary_Data

**Debayan Purkait, Farhana Islam, Padmaja P. Mishra**
**Chemical Sciences Division, Saha Institute of Nuclear Physics, India**
**Homi Bhaba National Institute (HBNI), India**

| Terminal PIFE Construct | Sequence (5' to 3') |
|---|---|
| **Biotin-DNA** | Biotin-AAAAAGTGTGTGTGT |
| **RecD2_Cy3_PIFE1** | /Cy3/TACAGCTACCTAGTCGATGTGCATACTACGGCACACACACAC |
| **RecD2_0** | GCCGTAGTATGCACATCGAC |

| Junction PIFE Construct | Sequence (5' to 3') |
|---|---|
| **Biotin-DNA** | Biotin-AAAAAGTGTGTGTGT |
| **RecD2_Cy3_PIFE2** | TACAGCTACCTA/Cy3/GTCGATGTGCATACTACGGCACACACACAC |
| **RecD2_0** | GCCGTAGTATGCACATCGAC |

| Internal PIFE Construct | Sequence (5' to 3') |
|---|---|
| **Biotin-DNA** | Biotin-AAAAAGTGTGTGTGT |
| **RecD2_Cy3_PIFE3** | TACAGCTACCTAGTCGATGTGCAT/Cy3/ACTACGGCACACACACAC |
| **RecD2_0** | GCCGTAGTATGCACATCGAC |

| Terminal FRET Construct | Sequence (5’ to 3’) |
|---|---|
| **Biotin-DNA** | Biotin-AAAAAGTGTGTGTGT |
| **RecD2_1_Cy3** | /Cy3/TACAGCTACCTAGTCGATGTGCATACTACGGCACACACACAC |
| **RecD2_1_Cy5** | GCCGTAGTATGCACATCGAC/Cy5/ |

| Junction FRET Construct | Sequence (5’ to 3’) |
|---|---|
| **Biotin-DNA** | Biotin-AAAAAGTGTGTGTGT |
| **RecD2_2_Cy3** | TACAGCTACCTA/Cy3/GTCGATGTGCATACTACGGCACACACACAC |
| **RecD2_1_Cy5** | GCCGTAGTATGCACATCGAC/Cy5/ |

| Internal FRET Construct | Sequence (5’ to 3’) |
|---|---|
| **Biotin-DNA** | Biotin-AAAAAGTGTGTGTGT |
| **RecD2_3_Cy3** | TACAGCTACCTAGTCGATGTGCAT/Cy3/ACTACGGCACACACACAC |
| **RecD2_2_Cy5** | GCCGTAGT/Cy5/ATGCACATCGAC |

**Table 1: Details of the individual ssDNA used in the experiments to make different working combinations of dsDNA.**

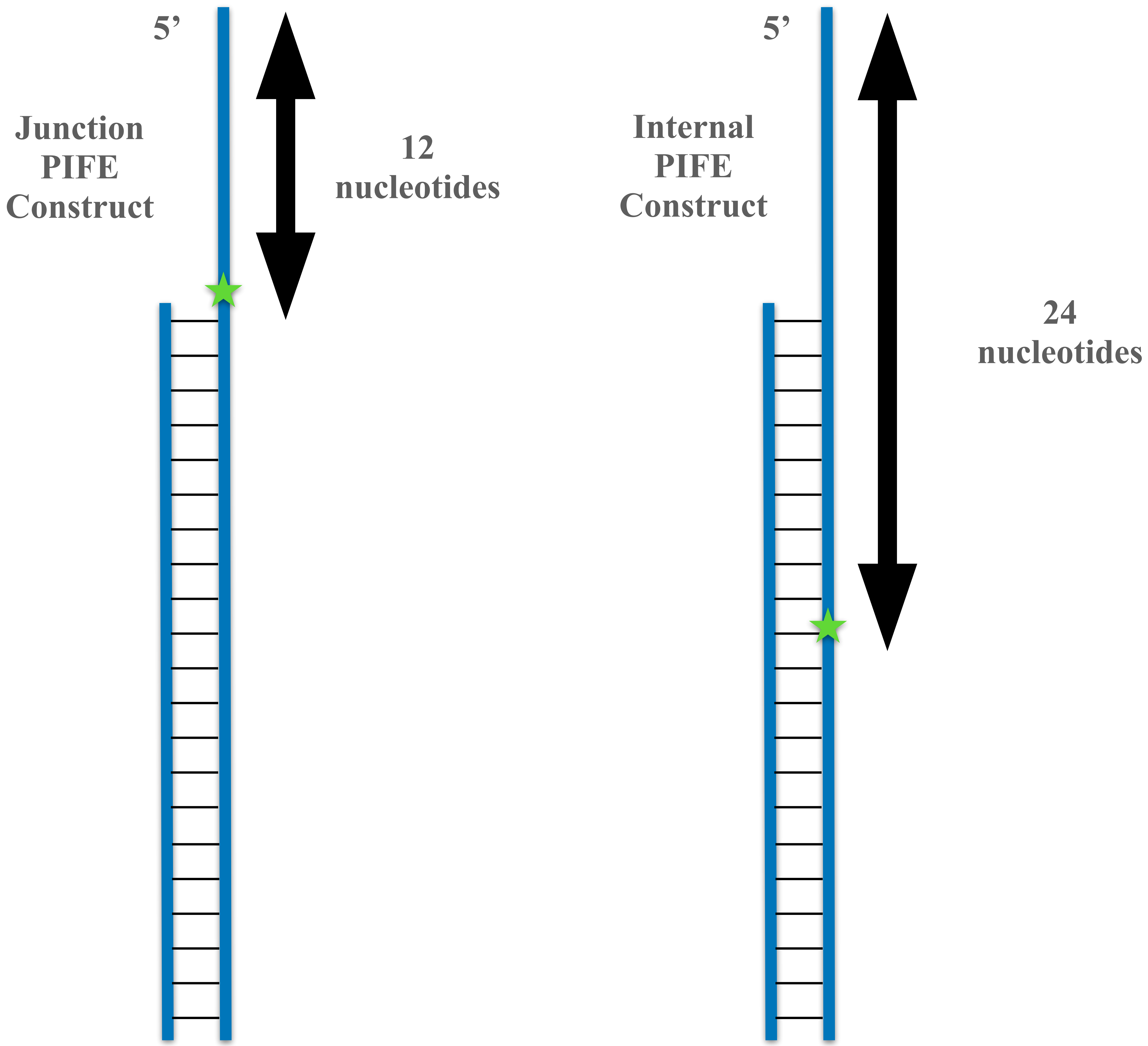


**Figure 1: Schematic representations of the Junction PIFE and Internal PIFE Construct**

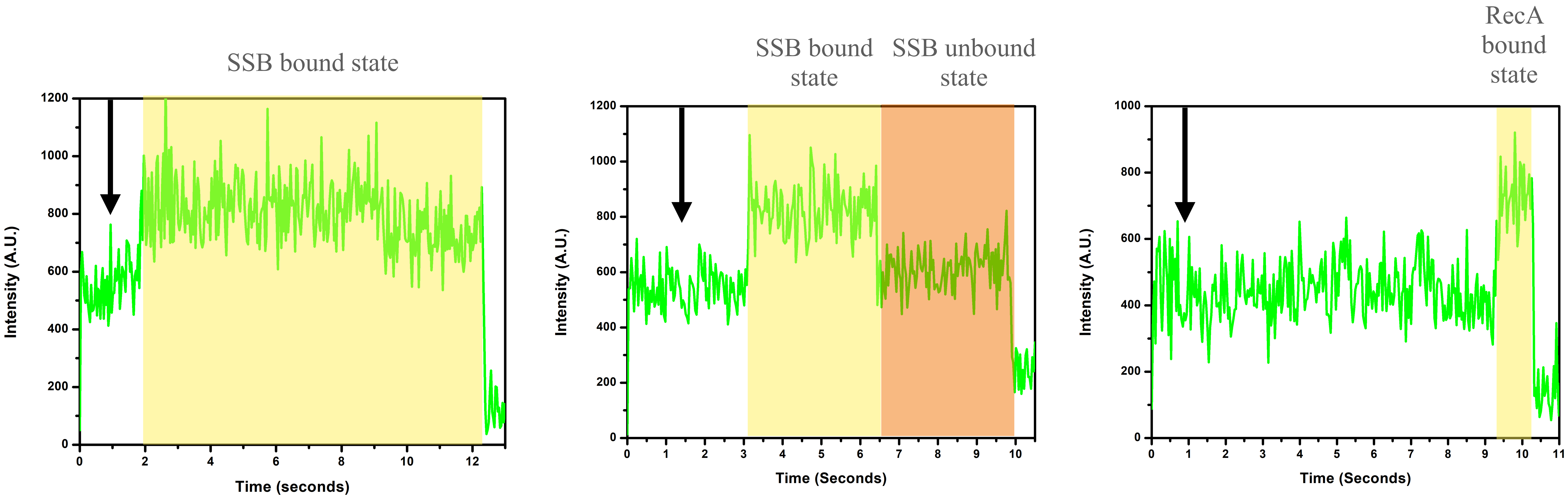


**Figure 2: smPIFE traces of binding events of SSB-homotetramer and RecA with the Terminal PIFE Construct. The black arrow indicates the time-point at which the proteins were introduced.**

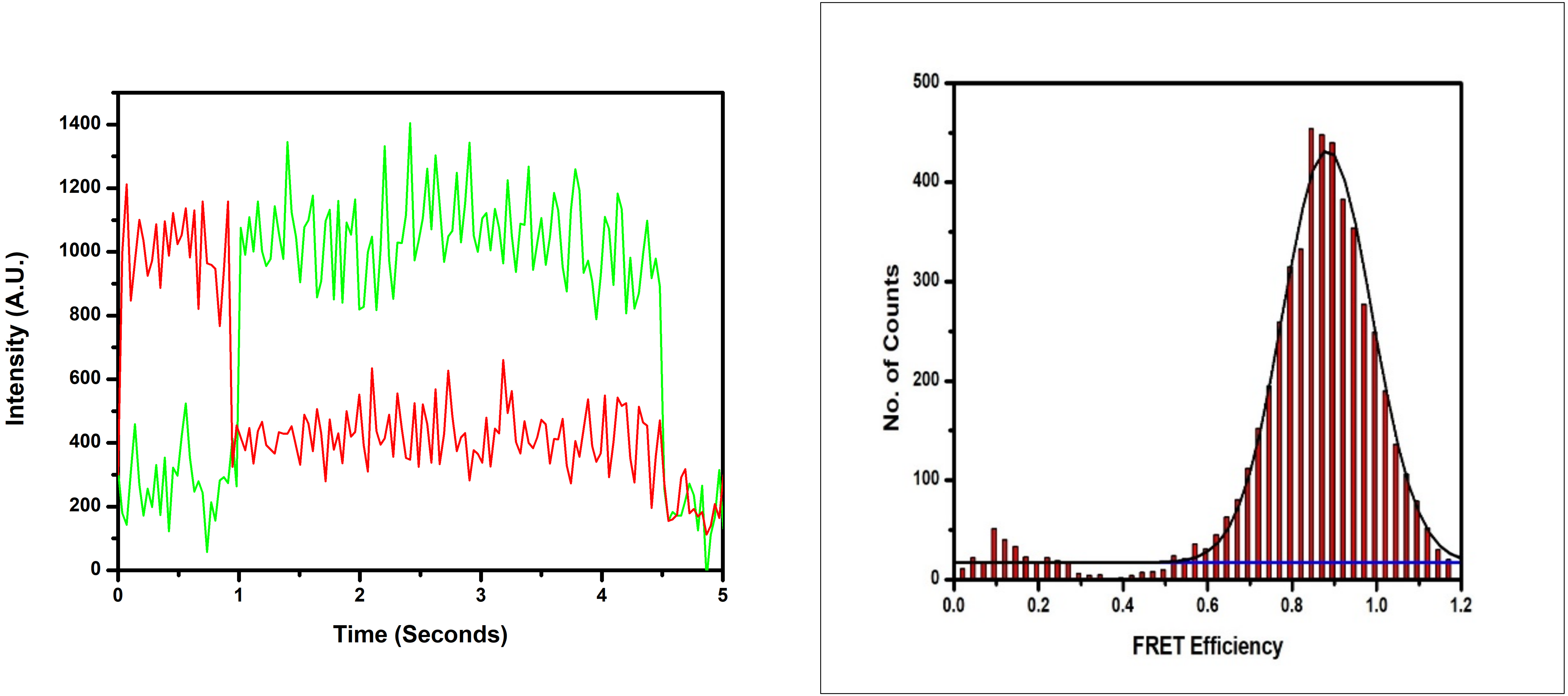


**Figure 3: smFRET trace and histogram of the Terminal FRET Construct. The presence of both low and high FRET state indicates the conformational dynamics of the 5' platform, however, the dominant High FRET population depicts that the hairpin is the stable form.**

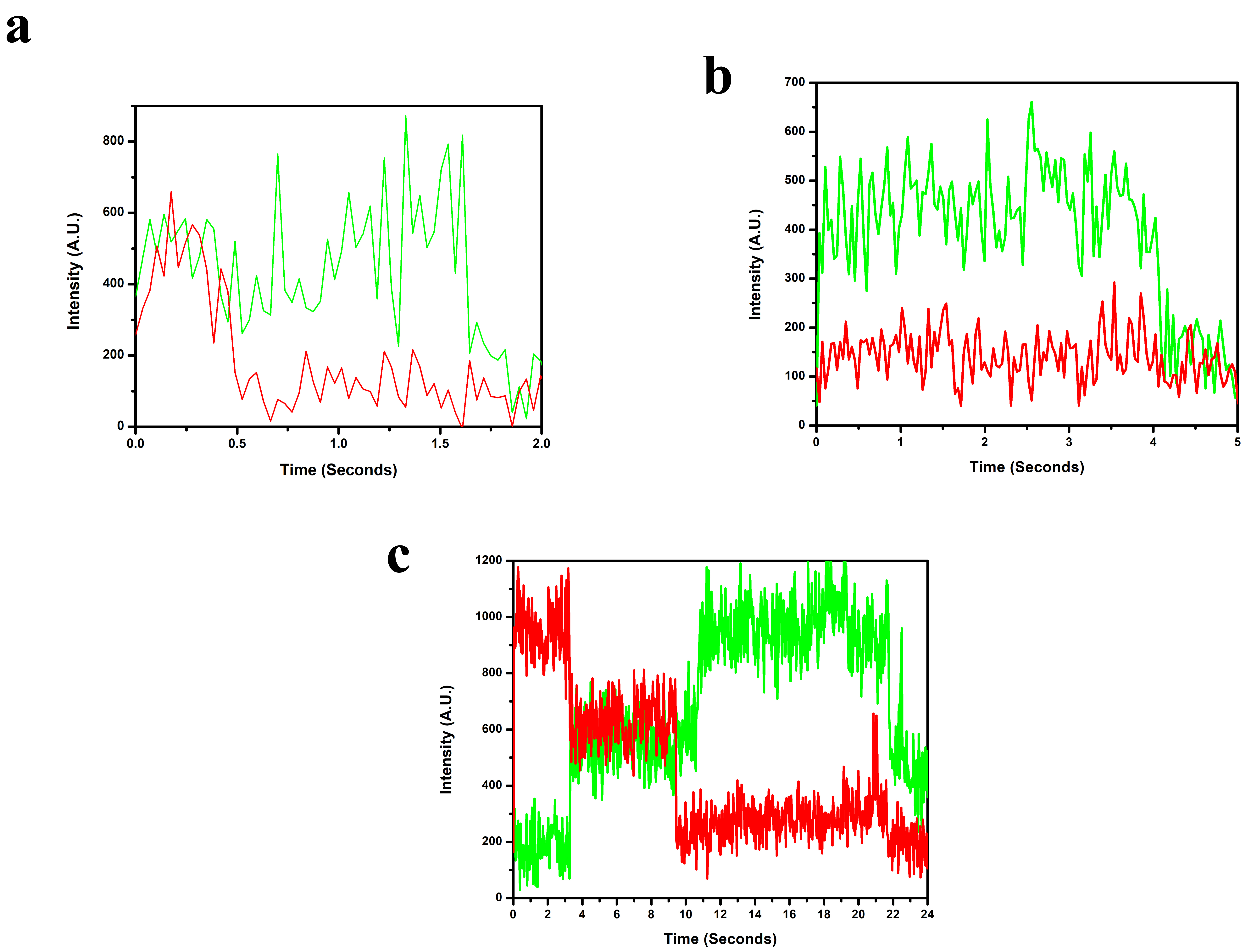


**Figure 4: smFRET traces of Terminal FRET Construct with (a) DrRecD2 (b) RecA & AMP-PNP (c) SSB**

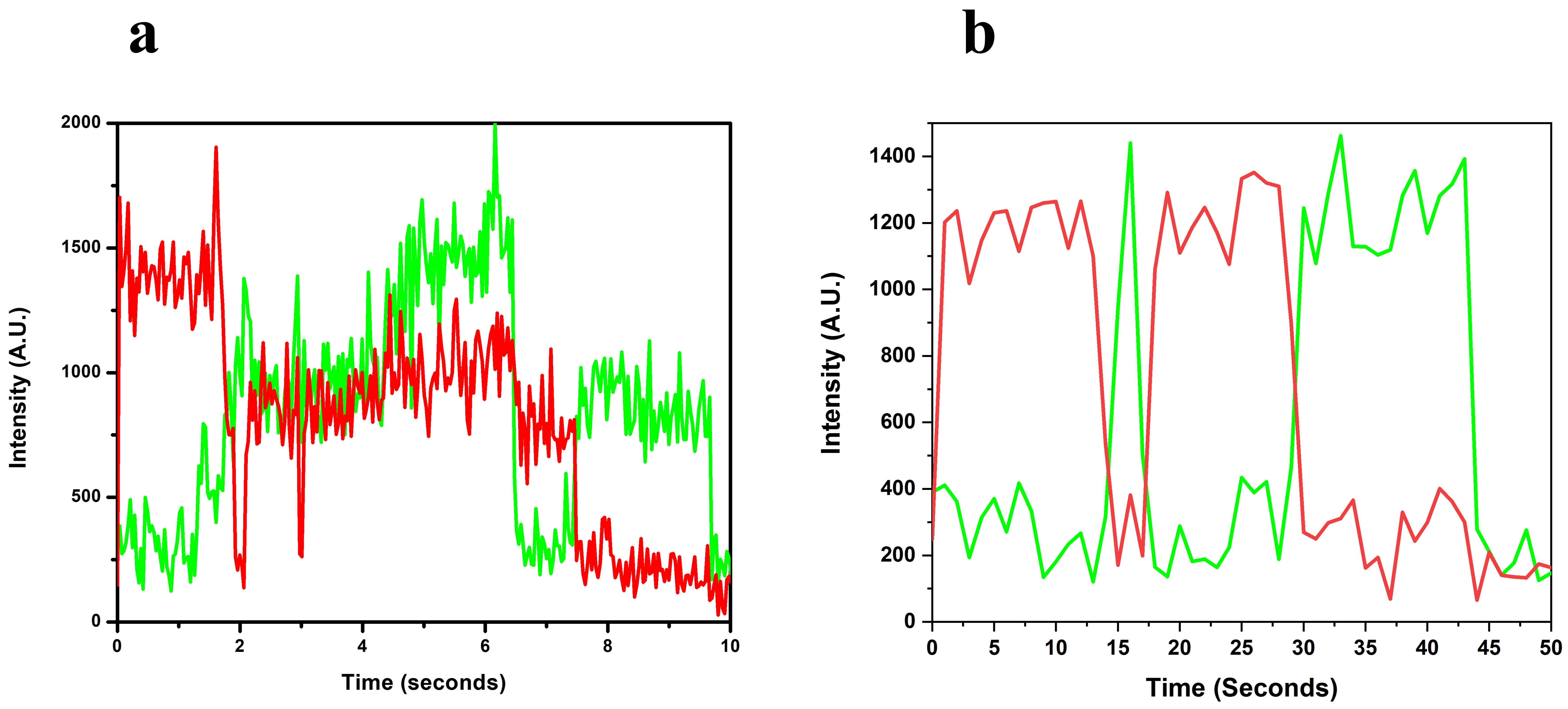


**Figure 5: smFRET traces of Terminal FRET Construct with (a) SSB & RecA & AMP-PNP (b) SSB & RecD2**

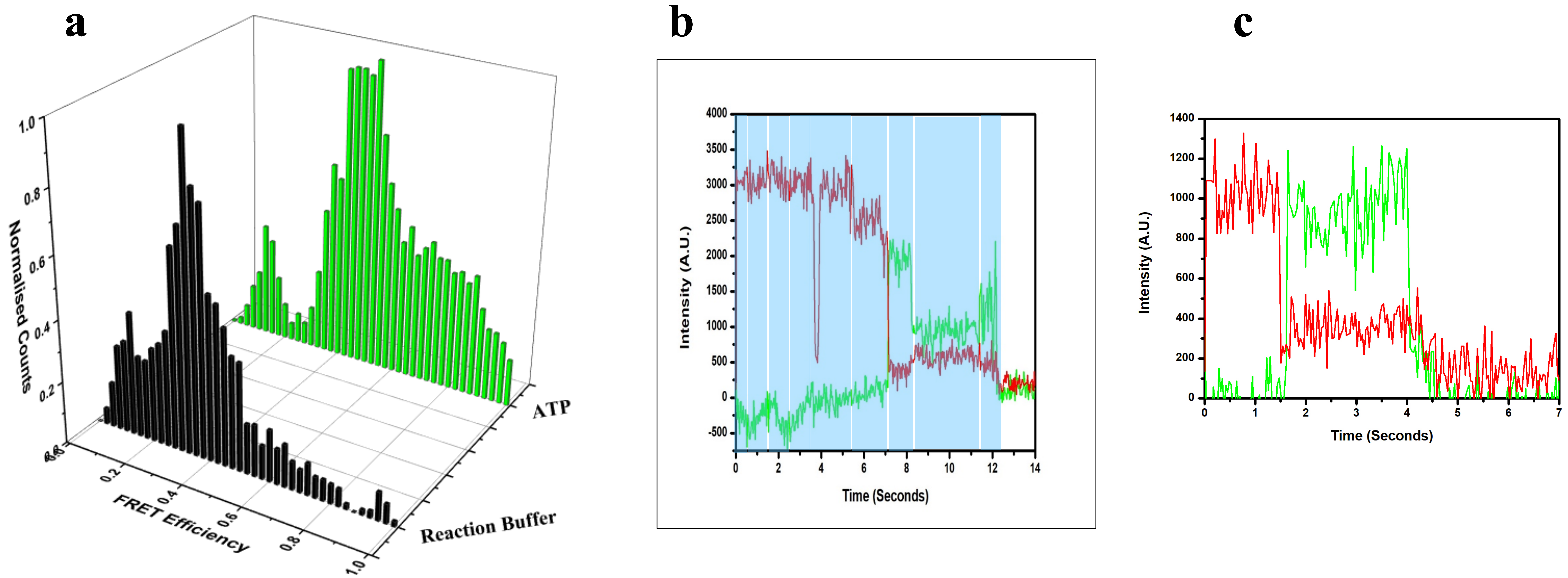


**Figure 6: (a) Comparative 3D histogram of FRET efficiencies and (b & c) smFRET traces of the Junction FRET construct for control experiments. (b) The smFRET trace shows the molecular fraying phenomenon at the ssDNA-dsDNA junction, the shaded area in Blue shows 9 different transient FRET state. (c) The fluctuations is getting stabilised in presence of ATP (5 mM)**

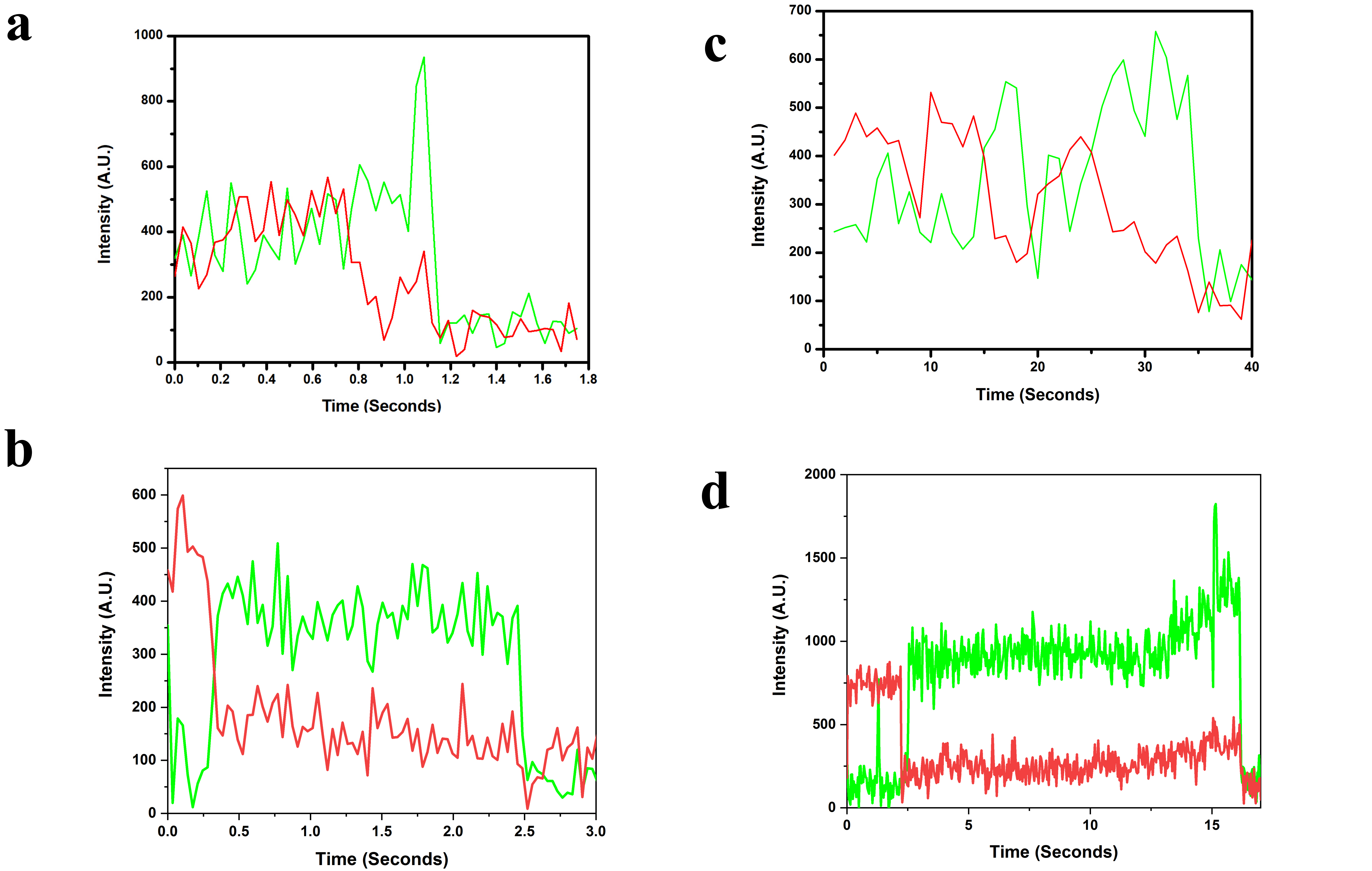


**Figure 7: smFRET traces of the Junction FRET Construct. (a) Only DrRecD2 (b) DrRecD2 & 5 mM AMP-PNP (c) DrRecD2 & 5 mM ATP (d) DrRecD2 & SSB & 5 mM ATP**

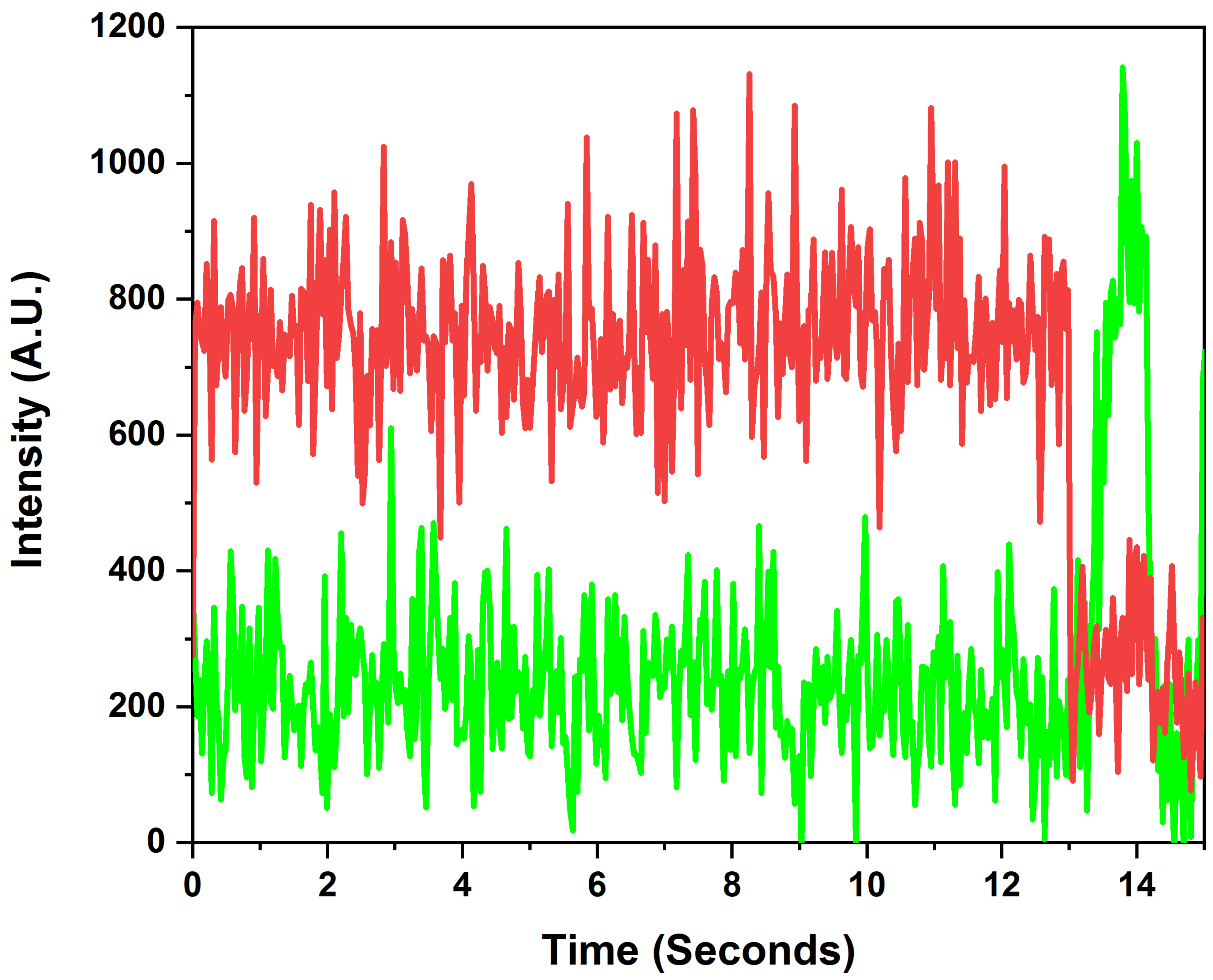


**Figure 8: smFRET traces of the Internal FRET Construct in Reaction Buffer**

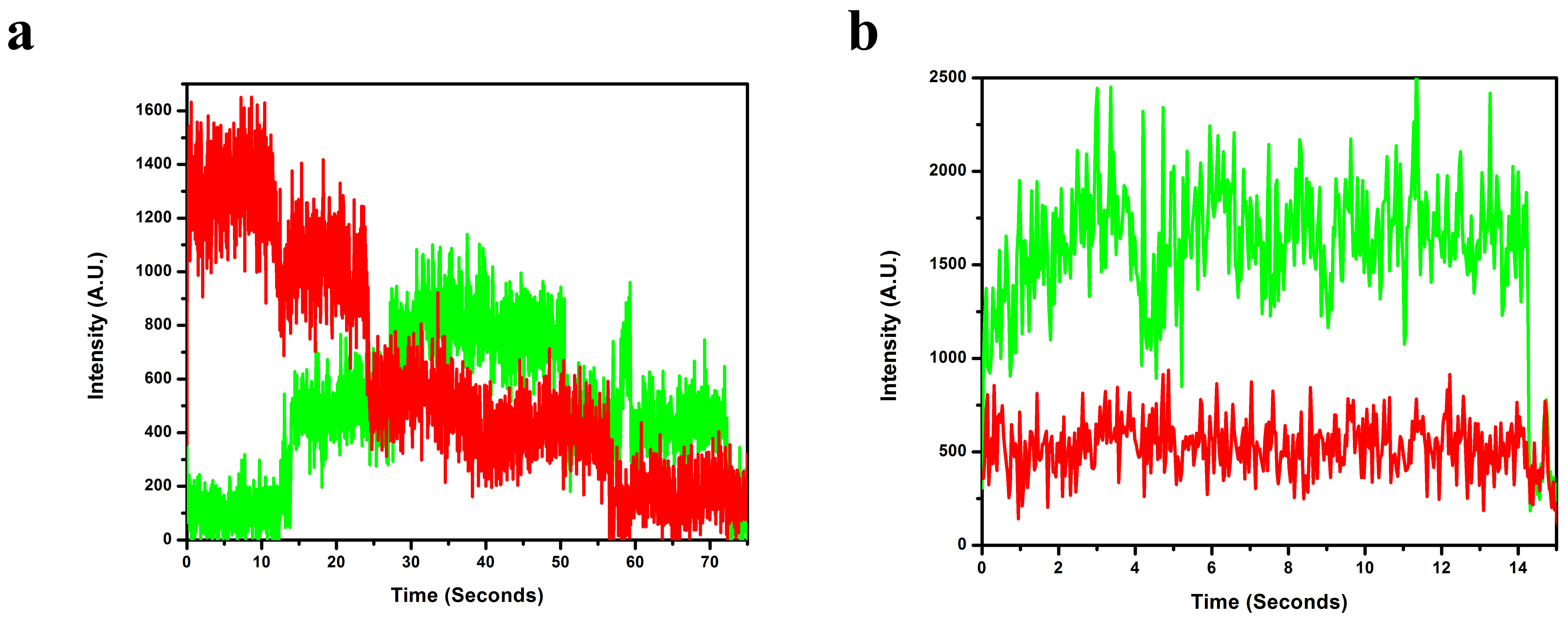


**Figure 9: smFRET traces of the Internal FRET Construct. (a) DrRecD2 & RecA & 5 mM AMP-PNP (b) DrRecD2 & RecA & 5 mM ATP**
